# Optimization of insect odorant receptor trafficking and functional expression via transient transfection in HEK293 cells

**DOI:** 10.1101/669127

**Authors:** Fabio Miazzi, Carolin Hoyer, Silke Sachse, Markus Knaden, Dieter Wicher, Bill S. Hansson, Sofia Lavista-Llanos

**Affiliations:** Max Planck Institute for Chemical Ecology, Department of Evolutionary Neuroethology, Hans-Knöll-Str. 8, 07745, Jena, Germany; Max Planck Institute for Chemical Ecology, Max Planck Research Group Predators and Prey, Hans-Knöll-Str. 8, 07745, Jena, Germany

**Author notes:** these authors share seniority (Bill S. Hansson, Sofia Lavista-Llanos). Correspondence to be sent to: Fabio Miazzi, Max Planck Institute for Chemical Ecology, Max Planck Research Group Predators and Prey, Hans-Knöll-Str. 8, 07745, Jena, Germany.

**Keywords:** *Drosophila melanogaster*, ORs, Rhodopsin, HCN1

## Abstract

Insect odorant receptors show a limited functional expression in various heterologous expression systems including insect and mammalian cells. This may be in part due to the absence of key components driving the release of these proteins from the endoplasmic reticulum and directing them to the plasma membrane. In order to mitigate this problem we took advantage of small export signals within the human HCN1 and Rhodopsin that have been shown to promote protein release from the endoplasmic reticulum and the trafficking of post-Golgi vesicles, respectively. Moreover, we designed a new vector based on a bidirectional expression cassette to drive the functional expression of the insect odorant receptor co-receptor (Orco) and an odor-binding odorant receptor, simultaneously. We show that this new method can be used to reliably express insect odorant receptors in HEK293 cells via transient transfection and that is highly suitable for downstream applications using automated and high-throughput imaging platforms.

## Introduction

Insect odorant receptors (ORs) are seven transmembrane-domain proteins, with an inverted topology compared to G protein-coupled receptors (GPCRs) (Benton et al., 2006; Butterwick et al., 2018), responsible for the detection of a vast number of chemically diverse odorants including pheromones (Hallem and Carlson, 2006; Touhara and Vosshall, 2009). They form heteromeric cation channels, constituted by an odor-binding receptor (OrX) and a highly conserved co-receptor named Orco (Krieger et al., 2003; Jones et al., 2005; Neuhaus et al., 2005; Benton et al., 2006; Sato et al., 2008; Wicher et al., 2008; Vosshall and Hansson, 2011). Several methods have been used to express insect ORs in heterologous systems in order to characterize their ligand specificity and to study their functional properties (Fleischer et al., 2018). For example, the so-called “empty neuron system” allows the ectopic expression of ORs in subsets of olfactory sensory neurons of the vinegar fly *Drosophila melanogaster* lacking the endogenous OrX, but with a fully functional Orco (Dobritsa et al., 2003; Kurtovic et al., 2007; Gonzalez et al., 2016). Moreover, *in vitro* expression systems allow the heterologous expression of OR proteins in animal cells including – among others – *Xenopus* oocytes (Nakagawa et al., 2005; Wang et al., 2010; Nakagawa and Touhara, 2013), insect cells (Kiely et al., 2007; Tsitoura et al., 2010; German et al., 2013) and mammalian cells (e.g. HEK293 and CHO) using vectors for transient expression (Sato et al., 2008; Wicher et al., 2008) or as stable lines (Große-Wilde et al., 2006; Jones et al., 2011; Corcoran et al., 2014).

Each of these methods offers several advantages, but also bears disadvantages: the “empty neuron system” allows the expression of OR proteins in an environment that is very similar to their native olfactory sensillum. However, the generation of transgenic fly lines and the electrophysiological recording tests can be time consuming and are not suitable for high-throughput screening experiments. *Xenopus* oocytes have been successfully used for extensive screenings of odors and are compatible with automated platforms, but the costs associated with *Xenopus* rearing or the purchase of oocytes can be prohibitive for very large screenings. Stable and inducible cell lines, on the other hand, originate from cell clones that successfully integrated in their genome an expression cassette containing a tetracycline-dependent promoter driving the expression of Orco and an odor-binding OrX. Such system represents a sort of “gold standard” as cell variability is strongly reduced, compared to transiently transfected cells, by selection of monoclonal populations and non-induced cells constitute an optimal internal negative control. Moreover, the selection of a monoclonal cell population represents – to date – the most effective approach to deal with the limited functional expression of ORs in the plasma membrane in heterologous systems due to an impaired intracellular trafficking of ORs both in insect (German et al., 2013) and mammalian cells (Halty-deLeon et al., 2016).

Although a thorough investigation of the bottlenecks in insect OR intracellular trafficking has not been performed yet, their retention within intracellular membranes (German et al., 2013) may be due to the lack of specialized components of the OR release mechanism from the endoplasmic reticulum (ER) and the Golgi apparatus in heterologous systems. Several membrane proteins have been shown to direct their intracellular trafficking through small peptide regions. Among others (Schülein et al., 1998; Ammon et al., 2002), it has been shown that an N-terminal peptide (^106^VNKFSL^111^) from the human HCN1 channel facilitates the exit of HCN1 proteins from the ER (Pan et al., 2015), and the C-terminal portion of the human Rhodopsin (^344^QVAPA^348^) ‒ containing the “VxPx” motif ‒ is sufficient to promote the formation of post-Golgi vesicles and their trafficking to the plasma membrane via microtubule-mediated transport (Tai et al., 1999; Deretic et al., 2005; Mazelova et al., 2009; Lodowski et al., 2013).

In this work, we investigated whether insect ORs tagged at their N-terminus with the HCN1 and Rhodopsin signal peptides show an enhanced trafficking to the plasma membrane in mammalian cells and reach satisfactory functional expression levels even after transient transfection without clonal selection. Moreover, in order to increase the efficiency of gene co-transfection, we designed a new vector based on a commercially available dual-promoter plasmid for mammalian transient transfection. In this way, we aimed at creating a new fast and inexpensive strategy to reliably express insect ORs in mammalian cells for a broad range of application, including automated and high-throughput imaging systems.

## Materials and Methods

### Constructs

A human codon-optimized version of *D. melanogaster* Or47a (hOr47a) tagged at the N-terminus with the peptide “QVAPAGKPIPNPLLGLDSTVNKFSL” coding for the human Rhodopsin ^344^QVAPA^348^ peptide (minimal Rho tag, abbreviated as "mRho" or "R" tag), a V5 tag (GKPIPNPLLGLDST) and the human HCN1 peptide ^106^VNKFSL^111^ (minimal ER export tag, abbreviated as "mER" or "E" tag) ‒ this construct is hereafter called mRho.V5.mER.hOr47a, or R.E.hOr47a ‒ was synthesized and subcloned into the pcDNA3.1(-) vector (Cat. Nr. V79520, Invitrogen, Carlsbad, CA) using the XbaI and XhoI restriction sites by Eurofins Genomics GmbH (Ebersberg, Germany). Constructs bearing only the V5 and mER peptides (V5.mER.hOr47a, or abbreviated E.hOr47a) and the V5 tag only (V5.hOr47a, or abbreviated hOr47a) were obtained from the mRho.V5.mER.hOr47a construct via PCR using the Advantage 2 PCR kit (Cat. Nr. 639206, Takara, Kusatsu, Japan) using the E.hOr47a_fwd and hOr47a_fwd forward primers, and the common (E.)hOr47a_rev reverse primer, respectively. A previously described non-human codon optimized version of *D. melanogaster* Orco cloned in pcDNA3.1(-) (Mukunda et al., 2014) was used for co-transfection of the hOr47a constructs in pcDNA3.1(-).

We created a high-copy number bidirectional expression vector (hence called pCMV-BI) by inserting the bidirectional promoter cassette flanked by the termination regions of the pBI-CMV1 vector (Cat. Nr. 631630 Clontech, Mountain View, CA), in the pCMVTNT (Cat. Nr. L5620, Promega, Madison, WI) vector backbone. Both regions were amplified using Phusion high-fidelity polymerase (Cat. Nr. M0530, New England Biolabs, Ipswich, MA) with the pBI-CMV1_fwd, pBI-CMV1_rev, and pCMVTNT_fwd, pCMVTNT_rev primer couples, respectively. Fragments were assembled using the NEBuilder HiFi DNA Assembly Master Mix (Cat. Nr. E2621, New England Biolabs).

A human codon-optimized version of *D. melanogaster* Orco (hOrco) tagged at the N-terminus with a myc tag (5’-GAACAGAAACTGATCTCTGAAGAAGACCTG-3’) was synthesized by Eurofins Genomics GmbH and subcloned into the pCMV-BI vector using the BamHI and HindIII restriction sites. A version of hOrco bearing the β-globin/IgG chimeric intron from the pCMVTNT vector within the hOrco sixth transmembrane domain was constructed by Phusion polymerase amplification using the hOrcoExon1_fwd, hOrcoExon1_rev, chimeric_intron_fwd, chimeric_intron_rev, hOrcoExon2_fwd and hOrcoExon2_rev primers. A new vector called pDmelOR was then created by cloning this intron-containing version of hOrco into the pCMV-BI vector, after digestion with BamHI and HindIII, using the NEBuilder HiFi DNA Assembly kit.

The mRho.V5.mER.hOr47a construct was inserted in the pDmelOR vector after linearization with EcoRI, using the BI-R.E.hOr47a_fwd and BI-R.E.hOr47a_rev using the NEBuilder HiFi DNA Assembly Master Mix (pDmelOR-mRho.V5.mER.hOr47a, or pDmelOR-R.E.hOr47a). Moreover, a human codon-optimized versions of *D. melanogaster* Or56a (hOr56a) was synthesized by Eurofins Genomics GmbH and inserted in the pDmelOR vector after linearization with EcoRI, together with the mRho.V5.mER tag using the following set of primers: hOr56a_fwd, hOr56a_rev, hOr56aTag_fwd, hOr56aTag_rev (pDmelOR-mRho.V5.mER.hOr56a, or pDmelOR-R.E.hOr56a).

All sequences were verified by Sanger sequencing (by Eurofins Genomics GmbH and the Department of Entomology, Max Planck Institute for Chemical Ecology, Jena). The CMV enhancer region of the pCMV-BI empty plasmid was sequenced using the pBI-CAG_for and pBI-CAG_rev primers. Sequencing of CMV promoter and termination regions from pCMV-BI and related constructs required a preliminary digestion with restriction enzymes, band isolation and purification after gel electrophoresis and possibly amplification of target sequences due to the presence of two very similar CMV promoter and SV40 polyA regions. The pCMV-BI empty plasmid was digested with DraIII-HF (Cat. Nr. R3510, New England Biolabs) and NcoI-HF (Cat. Nr. R3193, New England Biolabs). The pDmelOR vector was sequenced after digestion with XhoI and PvuI-HF (Cat. Nr. R3150, New England Biolabs); constructs with inserted Or47a and Or56a genes were cut with XhoI and NaeI (Cat. Nr. R0190, New England Biolabs). If required, the inserted Orco gene was subsequently amplified from the Orco-bearing purified plasmid fragment with the pBI-Orco_for and pBI-Orco_rev primers, and the tuning receptor gene with the pBI-OrX_fwd, pBI-OrX_rev primers using the Advantage 2 PCR kit and the resulting PCR products were sequenced. Primer sequences are listed in the Supplementary Table 1.

Full sequences for the reported constructs and plasmid availability information are accessible at Addgene (https://www.addgene.org) with the following reference numbers: pcDNA3.1(-)-mRho.V5.mER.hOr47a: #126472; pcDNA3.1(-)-V5.mER.hOr47a: #126473; pcDNA3.1(-)-V5.hOr47a: #126474; pCMV-BI: #126475; pDmelOR: #126476; pDmelOR-mRho.V5.mER.hOr47a: #126478; pDmelOR-mRho.V5.mER.hOr56a: #126479.

### Transient expression in mammalian cells

HEK293 cells (DSMZ no. ACC 305) were purchased from the Leibniz Institute DSMZ GmbH (Braunschweig, Germany) and grown in DMEM/F12 1:1 medium (Cat. Nr. 11320, Gibco, Life Technologies, Grand Island, NY, USA) supplied with 10% Fetal Bovine Serum at 37°C and 5% CO_2_. Cell transfection was performed by electroporation with an Amaxa 4D-Nucleofector (Lonza GmbH, Cologne, Germany) and the SF Cell Line 4D-Nucleofector X Kit (Lonza GmbH) using 80-90% confluent cells after dissociation by trypsinization. For each experiment (n = 1) shown in Figure 1 and Figure 2, 1.8 × 10^6^ cells were mixed with 0.6 µg of pcDNA3.1(-) bearing one of the tested Or47a constructs (or empty vector), and 0.6 µg of pcDNA3.1(-) bearing Orco (or empty vector), and then loaded in an electroporation cuvette (total of 1.2 µg/cuvette plasmid DNA). For experiments shown in Figure 2 with the pDmelOR-R.E.hOr47a construct, 0.8 µg of plasmid per cuvette was used (total of 0.8 µg/cuvette plasmid DNA). For each experiment (n = 1) shown in Figure 3, two cuvettes with ∼1.1 × 10^6^cells/cuvette were mixed with 0.8 µg/cuvette of pDmelOR-R.E.hOr47a, or two cuvettes with ∼0.67 × 10^6^ cells/cuvette were transfected with 0.8 µg/cuvette of pDmelOR-R.E.hOr56a; for control experiments with the pCMV-BI plasmid two cuvettes with ∼1 × 10^6^ cells/cuvette were transfected with 0.8 µg/cuvette of pCMV-BI. Electroporated cells were cultured on poly-l-lysine (0.01%, Sigma-Aldrich, Steinheim, Germany) coated coverslips at ∼3 × 10^5^cells/well (6 wells per cuvette) in a 24 wells plate (for experiments shown in Figure 1 and Figure 2), or they were split in 48 wells in Poly-D-Lysine Cellware 96-well black/clear plate (BD Biosciences, San Jose, CA, Cat. Nr. 354640) (for experiments shown in Figure 3), with a 1:1 mixture of DMEM zero Ca^2+^ (Gibco, Cat. Nr. 21068) supplemented with 1% Roti-Cell glutamine solution (Cat. Nr. 9183.1, Carl Roth), and F12 (Gibco, Cat. Nr. 21765) in a 1:1 mixture, supplemented with 10% Fetal Bovine Serum. Zero Ca^2+^ DMEM was preferred to standard DMEM in a 1:1 mix with F12, as we previously showed that such low-Ca^2+^ culture medium better supports the functional expression of insect ORs in HEK293 cells (Miazzi et al., 2019).

**Figure 1.**
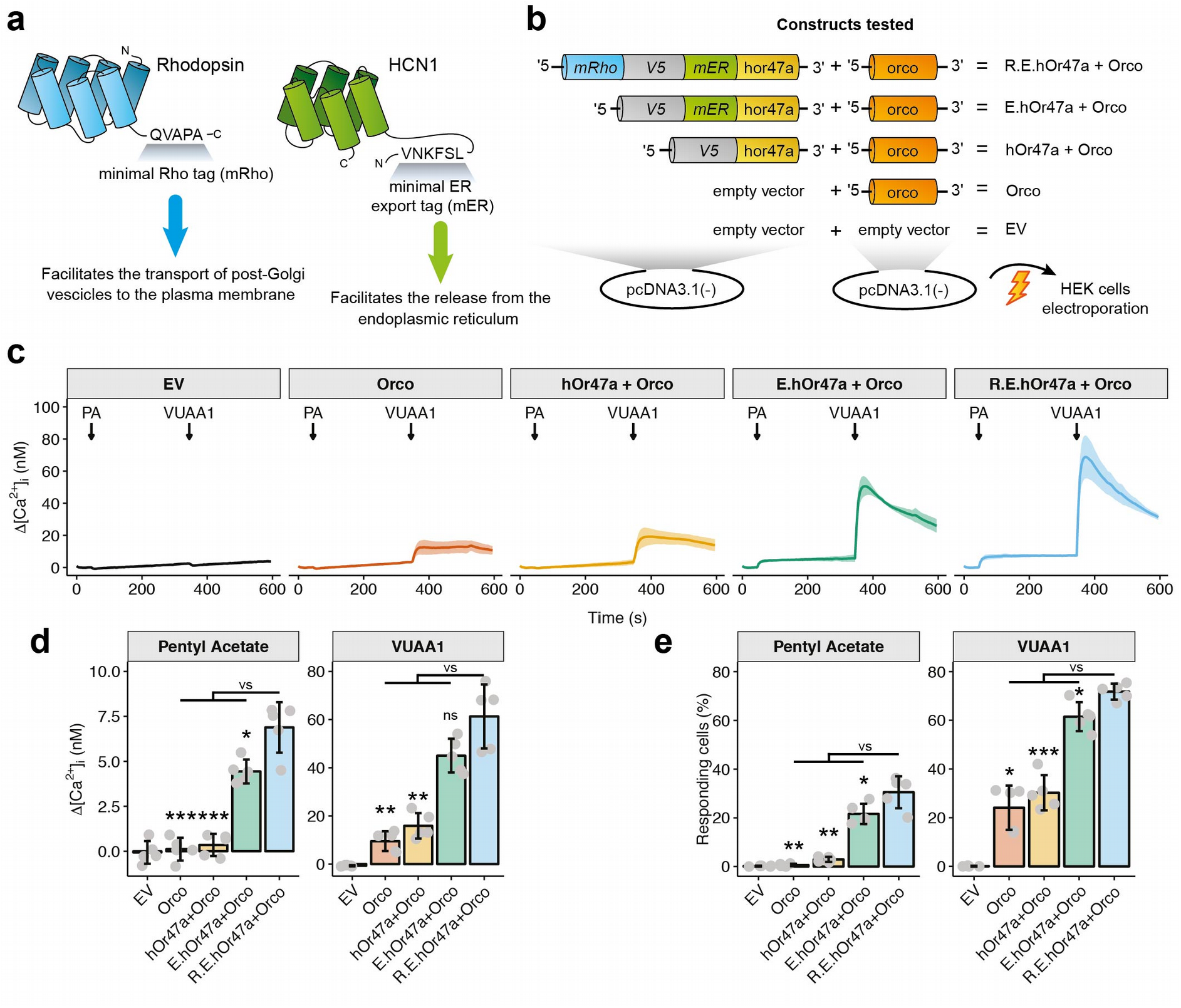
Optimization of OR trafficking to the plasma membrane in HEK293 cells. (a) The human HCN1 receptor sequence ^106^VNKFSL^111^ encoding a minimal endoplasmic reticulum release signal ("mER", or "E") and the human Rhodopsin sequence ^344^QVAPA^348^ encoding a minimal Rho tag ("mRho", or "R") were used to enhance the functional expression of insect ORs by promoting their exit from the endoplasmic reticulum and the transport of post-Golgi vesicles to the plasma membrane, respectively. (b) Schematic representation of the constructs tested. HEK293 cells were co-transfected by electroporation with two pcDNA3.1(-) plasmid constructs. The first, with the insertion of a human codon-optimized version of the *D. melanogaster* Or47a (hOr47a) tagged at the N-terminus with: a peptide composed of the mRho, mER and a V5 tag – to detect the receptor via immunochemistry if necessary – (R.E.hOr47a construct), or tagged only with the mER and V5 peptides (E.hOr47a) or the V5 tag alone (hOr47a). The second construct was constituted by a non codon-optimized version of *D. melanogaster* Orco co-receptor. As controls, cells were co-transfected with the Orco co-receptor together with the empty vector backbone (EV) instead of the hOr47a construct, or with the empty vector alone. (c) Changes in intracellular calcium concentration over time (Δ[Ca^2+^]_i_) in HEK293 cells transfected with the constructs shown in panel (b), following stimulations with 100 µl of 100 µM of the Or47a agonist pentyl acetate (PA) at 50 s and the synthetic Orco agonist VUAA1 at 350 s. Graphs represent mean ± SD. n = 5 for each panel. Each n = 1 represents the mean of all imaged cells coming from each independent electroporated cuvette (see Methods). (d) Intensity of calcium responses following a pentyl acetate or a VUAA1 stimulation in cells transfected with the constructs described in panel (b). Values were extracted 50 s after a stimulation with 100 µl of 100 µM pentyl acetate, and 20 s after a stimulation with 100 µl of 100 µM VUAA1. Mean ± SD values for pentyl acetate: EV, −0.07 ± 0.63; Orco, 0.11 ± 0.64; hOr47a + Orco, 0.35 ± 0.61; E.hOr47a + Orco, 4.44 ± 0.66; R.E.hOr47a + Orco, 6.89 ± 1.41. Statistical analysis for pentyl acetate: R.E.hOr47a + Orco versus (vs.) E.hOr47a + Orco: t = 3.52, p = 0.014; R.E.hOr47a + Orco vs. hOr47a + Orco: t = 9.52, p < 0.001; R.E.hOr47a + Orco vs. Orco: t = 9.81, p < 0.001. Mean ± SD values for VUAA1: EV, −0.68 ± 0.24; Orco, 9.52 ± 4.11; hOr47a + Orco, 15.88 ± 5.29; E.hOr47a + Orco, 45.02 ± 7.02; R.E.hOr47a + Orco, 61.32 ± 13.29. Statistical analysis for VUAA1: R.E.hOr47a + Orco versus (vs.) E.hOr47a + Orco: t = 2.42, p = 0.051; R.E.hOr47a + Orco vs. hOr47a + Orco: t = 7.10, p = 0.0015; R.E.hOr47a + Orco vs. Orco: t = 8.32, p = 0.0015. (e) Percentage of responding ROIs following a pentyl acetate or a VUAA1 stimulation. Mean ± SD values for pentyl acetate: EV, 0.18 ± 0.16; Orco, 0.64 ± 0.50; hOr47a + Orco, 2.90 ± 1.05; E.hOr47a + Orco, 21.61 ± 4.16; R.E.hOr47a + Orco, 30.51 ± 6.56. Statistical analysis for pentyl acetate: R.E.hOr47a + Orco versus (vs.) E.hOr47a + Orco: t = 2.56, p = 0.039; R.E.hOr47a + Orco vs. hOr47a + Orco: t = 9.29, p = 0.0015; R.E.hOr47a + Orco vs. Orco: t = 10.16, p = 0.0015. Mean ± SD values for VUAA1: EV, 0.06 ± 0.14; Orco, 24.12 ± 9.10; hOr47a + Orco, 30.24 ± 7.21; E.hOr47a + Orco, 61.49± 5.95; R.E.hOr47a + Orco, 71.76 ± 3.29. Statistical analysis for VUAA1: R.E.hOr47a + Orco versus (vs.) E.hOr47a + Orco: t = 3.38, p = 0.015; R.E.hOr47a + Orco vs. hOr47a + Orco: t = 11.71, p < 0.001; R.E.hOr47a + Orco vs. Orco: W = 25, p = 0.016 (Wilcoxon rank test). Unless otherwise stated, tests are unpaired two-tails Welch’t t-test. P values corrected for multiple comparisons using Holm’s correction. Bar plots represent mean ± SD, n = 5 for each treatment. * p < 0.05, ** p < 0.01, *** p < 0.001, ns = not significant.

**Figure 2.**
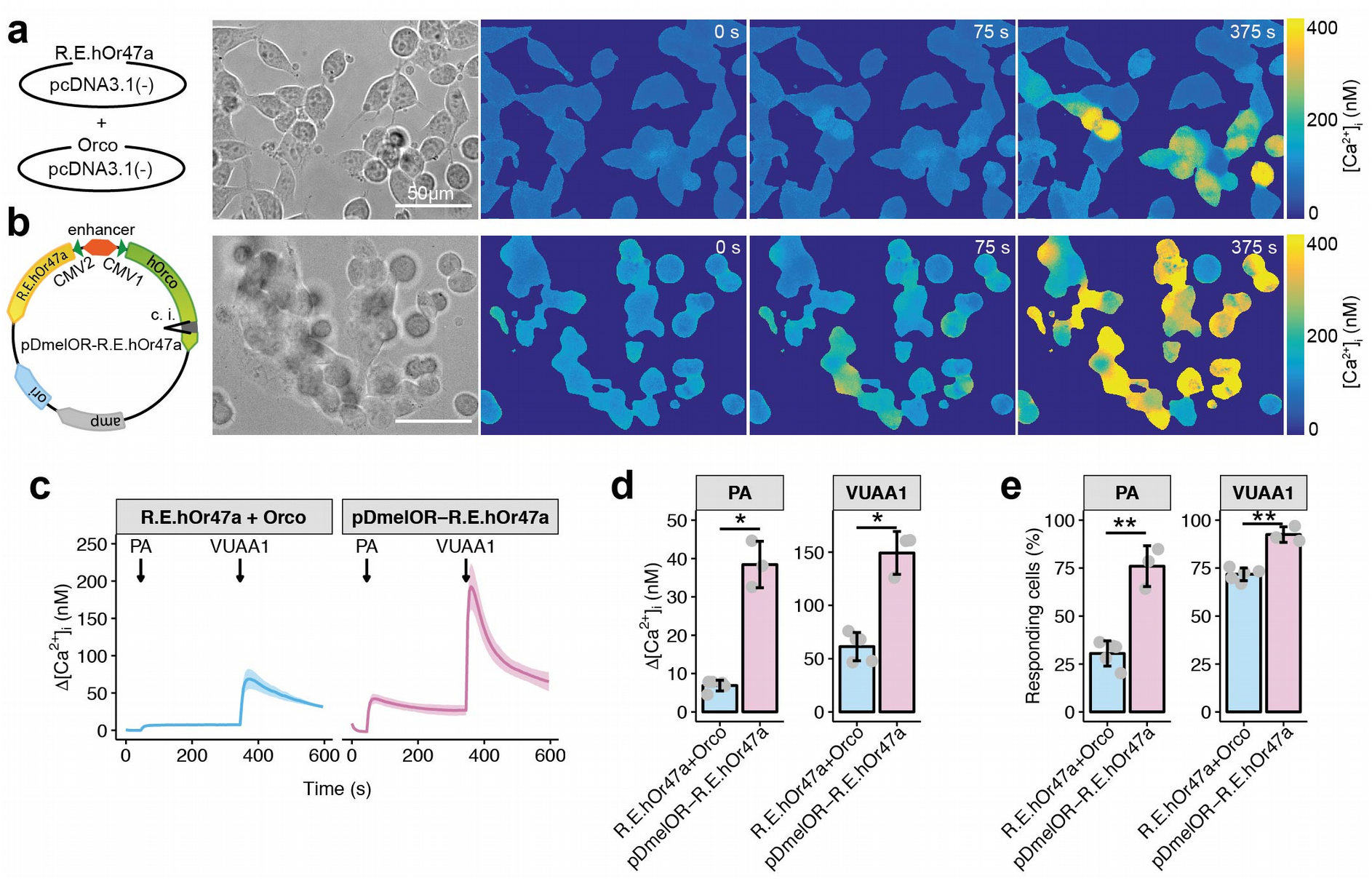
OR co-transfection via a bidirectional expression vector in HEK293 cells. (a-b) Examples of calcium imaging experiments with HEK293 cells co-transfected with the R.E.hOr47a + Orco constructs in two different pcDNA3.1(-) vectors (a) and with the pDmelOR-R.E.hOr47a housing both the hOr47a and a codon-optimized version of Orco (hOrco) bearing the β-globin/IgG chimeric intron (c.i.) (b). Panels represent – from left to right – a schematic representation of the constructs tested; transmission light image of the tested HEK293 cells; base level [Ca^2+^]_i_ of transfected HEK293 cells (0 s); [Ca^2+^]_i_ 25 s after application of 100 µl of 100 µM pentyl acetate (75 s) and 25 s after application of 100 µl of 100 µM VUAA1 (375 s). Scale bar = 50 µm. (c) Changes in intracellular calcium concentration over time (Δ[Ca^2+^]_i_) in HEK293 cells transfected with the construct shown in panels (a-b), following stimulations with 100 µl of 100 µM of the Or47a agonist pentyl acetate (PA) at 50 s and the synthetic Orco agonist VUAA1 at 350 s. Graphs represent mean ± SD. n = 5 for R.E.hOr47a + Orco and n = 3 for pDmelOR-R.E.hOr47a. Each n = 1 represents the mean of all imaged cells coming from independently electroporated cuvettes (see Methods). (d) Intensity of calcium responses following a pentyl acetate or a VUAA1 stimulation in cells transfected with the constructs described in panels (a-b). Δ[Ca^2+^]_i_ values were extracted 50 s after a stimulation with 100 µl of 100 µM pentyl acetate, and 20 s after a stimulation with 100 µl of 100 µM VUAA1. Mean ± SD values for pentyl acetate: R.E.hOr47a + Orco, 6.89 ± 1.41; pDmelOR-R.E.hOr47a, 38.43 ± 6.04. Statistical analysis for pentyl acetate: pDmelOR-R.E.hOr47a versus R.E.hOr47a + Orco, t = 8.90, p = 0.010, Two tailed Welch’s t test. Mean ± SD values for VUAA1: R.E.hOr47a + Orco, 61.32 ± 13.29; pDmelOR-R.E.hOr47a + Orco, 149.24 ± 20.12. Statistical analysis for VUAA1: pDmelOR-R.E.hOr47a versus R.E.hOr47a + Orco, W = 15, p = 0.036, Wilcoxon rank sum test. Graphs represent mean ± SD. n = 5 for R.E.hOr47a + Orco and n = 3 for pDmelOR-R.E.hOr47a. (e) Percentage of responding ROIs following a pentyl acetate or a VUAA1 stimulation. Mean ± SD values for pentyl acetate: R.E.hOr47a + Orco, 30.51 ± 6.56; pDmelOR-R.E.hOr47a, 76.01± 10.64. Statistical analysis for pentyl acetate: pDmelOR-R.E.hOr47a versus R.E.hOr47a + Orco: t = 6.68, p = 0.0073. Mean ± SD values for VUAA1: R.E.hOr47a + Orco, 71.76 ± 3.29; pDmelOR-R.E.hOr47a, 92.47± 4.03. Statistical analysis for VUAA1: pDmelOR-R.E.hOr47a versus R.E.hOr47a + Orco: t = 7.52, p = 0.0024. Two-tailed Welch’s t tests. Graphs represent mean ± SD. n = 5 for R.E.hOr47a + Orco and n = 3 for pDmelOR-R.E.hOr47a.

**Figure 3.**
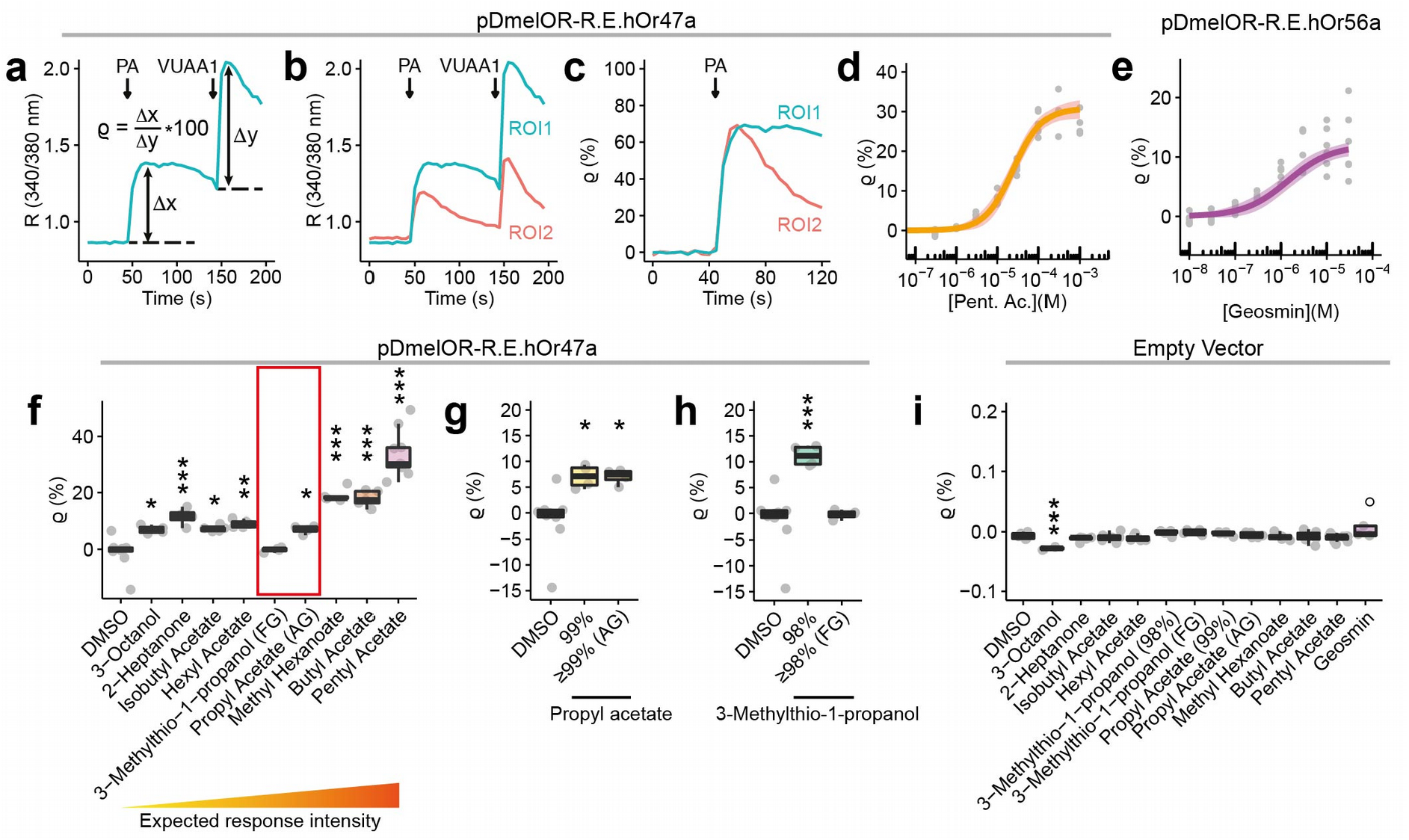
Analysis of insect OR agonist sensitivity and specificity by means of an automated imaging platform. (a) Normalization of the odor response respect to the VUAA1 internal control. After calculating the ratio (R) between the emission light at 340 nm and 380 nm, base levels were subtracted from peak agonist responses to obtain the odor, e.g. pentyl acetate (PA), and the VUAA1 net responses (∆x and ∆y, respectively). The ratio ϱ (*rho*) between the odor and VUAA1 net responses expressed in percentage (ϱ = [∆x/∆y]*100) can be used to account for the cell-specific OR expression level due to the transient transfection protocol. (b-c) Example showing how normalization of the odor response to the VUAA1 internal control can reduce response variability. (b) Time series of R (340/380 nm) ratios for two regions of interests (ROI1 and ROI2) showing different response intensities to the same odor and VUAA1 stimulation. (c) Time series of normalized ϱ values for the odor responses of the same ROI1 and ROI2 as shown in panel (b). ϱ values reveal that both ROI1 and ROI2 show a very similar maximal response to the PA stimulation. (d-e) Dose-response curve for HEK293 cells transfected with pDmelOR-R.E.hOr47a and stimulated with pentyl acetate (d), or with pDmelOR-R.E.hOr56a and stimulated with geosmin (e). Data were fitted to a 3-parameter logistic function using the drc package in R (see Supplementary Code). Values of the data fit (mean ± st. error) for pDmelOR-R.E.hOr47a: slope, −2.89 ± 0.42; upper limit, 30.80 ± 1.28; EC_50_ (log), −4.61 ± 0.06. EC_50_ values expressed in Molar (estimate, 2.5% and 97.5% confidence intervals): 2.45⋅10^−5^ (1.83⋅10^−5^; 3.29⋅10^−5^). Values of the data fit for pDmelOR-R.E.hOr56a: slope, −2.46 ± 0.58; upper limit, 11.93 ± 0.92; EC_50_ (log), −6.14 ± 0.13. EC_50_ values expressed in Molar (estimate, 2.5% and 97.5% confidence intervals): 7.19⋅10^−7^ (3.97⋅10^−7^; 1.30⋅10^−6^). Plot shows estimate ± S.E. For 5 ≤ n ≤ 6 (b) and n = 7 (c) for each odor concentration. (f) Odor response profile for Or47a using the pDmelOR-R.E.hOr47a construct. Odors are disposed according to their expected potency as agonists according to the DoOR database (see Supplementary Figure 2). Kruskall-Wallis rank sum test: χ^2^ = 50.148, df = 9, p < 0.001. Post-hoc Dunnett’s test for comparing each treatment versus (vs.) DMSO. 3-octanol vs. DMSO, p = 0.039; 2-heptanone vs. DMSO, p < 0.001; isobutyl acetate vs. DMSO, p = 0.013; hexyl acetate vs. DMSO, p = 0.002; 3-methylthio-1-propanol (FG) vs. DMSO, p = 1; propyl acetate (AG) vs. DMSO, p = 0.036; methyl hexanoate vs. DMSO, p < 0.001; butyl hexanoate vs. DMSO, p < 0.001; pentyl acetate vs. DMSO, p < 0.001. 4 ≤ n ≤ 10. The red box highlights two odors whose response intensities do not follow the expected trend, i.e. 3-methilthio-1-propanol and pentyl acetate. (g) The effect of propyl acetate is consistent across commercial stocks with different purity. The response to pentyl acetate was significant compared to the DMSO control when working solutions were prepared from a 99% purity stock odor (Sigma-Aldrich, Cat. Nr. 133108), and a ≥99% analytical standard grade (AG) stock odor (Fluka, Cat. Nr. 40858). Kruskall-Wallis rank sum test: χ^2^ = 10.088, df = 2, p = 0.0064. Post-hoc Dunnett’s test for comparing each treatment versus (vs.) DMSO. 99% vs. DMSO, p = 0.0147; ≥99% AG vs. DMSO, p = 0.0147. 4 ≤ n ≤ 9. (h) The effect of 3-methilthio-1-propanol is not consistent across commercial stocks with different purity. Responses were significant compared to the DMSO control when working solutions were prepared from a 98% purity stock odor (Sigma-Aldrich, Cat. Nr. 318396), but not from a ≥98% food/pharmaceutical (FG) grade stock odor (Sigma-Aldrich, Cat. Nr. W341509). Kruskall-Wallis rank sum test: χ^2^ = 8.681, df = 2, p = 0.013. Post-hoc Dunnett’s test for comparing each treatment versus (vs.) DMSO. 98% vs. DMSO, p = 0.904; ≥98% (FG) vs. DMSO: p < 0.001. 4 ≤ n ≤ 9. (i) Responses of HEK293 cells transfected with the pCMV-BI empty vector to odor stimuli (300 µM for all odors, except 30 µM for geosmin) and DMSO. Kruskall-Wallis rank sum test: χ^2^ = 27.115, df = 12, p = 0.0074. Post-hoc Dunnett’s test for comparing each treatment versus (vs.) DMSO. 3-octanol vs. DMSO, p = 0.00064; 2-heptanone vs. DMSO, p = 0.734; isobutyl acetate versus (vs.) DMSO, p = 0.996; hexyl acetate vs. DMSO, p = 0.949; 3-methylthio-1-propanol (98%) vs. DMSO, p = 0.762; 3-methylthio-1-propanol (FG) vs. DMSO, p = 0.572; propyl acetate (99%) vs. DMSO, p = 0.922; propyl acetate (AG) vs. DMSO, p > 0.999; methyl hexanoate vs. DMSO, p > 0.999; butyl acetate vs. DMSO, p > 0.999; pentyl acetate vs. DMSO, p = 0.961; geosmin vs. DMSO, p = 0.749. For geosmin, one data point (black circle in the bar plot) was considered as an outlier and was omitted from analysis. 2 ≤ n ≤ 11. *** p < 0.001, ** p < 0.01, * p < 0.05.

### Functional imaging

Cells were imaged 24 hours post-electroporation. Cells were incubated in Opti-MEM medium (Cat. Nr. 31985, Gibco), containing 5 µM Fura-2 acetoxymethyl ester (Molecular Probes, Invitrogen) for 30 min at room temperature. After incubation, cells were washed three times and kept throughout the experiment in standard extracellular solution (SES) containing 135 mM NaCl, 5 mM KCl, 1 mM CaCl_2_, 1 mM MgCl_2_, 10 mM HEPES, 10 mM glucose (pH = 7.4; osmolarity = 295 mOsmol/l). Excitation with 340 and 380 nm light for 150 ms per frame was obtained using a monochromator (Polychrome V, Till Photonics, GrYfelfing, Germany), coupled to an epifluorescence microscope (Axioskop FS, Zeiss, Jena, Germany) by means of a water immersion objective (LUMPFL40×W/IR/0.8; Olympus, Hamburg, Germany) and controlled by an imaging control unit (ICU, Till Photonics). Emitted light was separated by a 400 nm dichroic mirror, filtered with a 420 nm long-pass filter and acquired by a cooled CCD camera (Sensicam, PCO Imaging, Kelheim, Germany) controlled by the TILLVision 4.5 software (TILL Photonics). The final image resolution was 640 × 480 pixels in a frame of 175 × 130 µm. Experiments lasted 10 min with a sampling interval of 5 s. Stimuli consisting of 100 µl of pentyl acetate and VUAA1 solutions (100 µM in SES) were delivered via pipette in proximity of the objective at 50 and 350 seconds (s), respectively. Movies of the imaging recordings were exported as uncompressed TIFF files from the TILLVision software and analyzed using custom scripts for Fiji ImageJ2.0 (Schindelin et al., 2012; Rueden et al., 2017) (see Supplementary Code). Briefly, the free intracellular Ca^2+^ concentration ([Ca^2+^]_i_) was calculated after correcting for background, flat-field and movement (performed using the image stabilizer plugin for ImageJ ‒ http://www.cs.cmu.edu/~kangli/code/Image_Stabilizer.html) artifacts. [Ca^2+^]_i_ was calculated according to the equation:

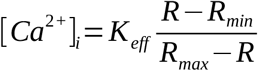

Where K_eff_ = 1747 nM, R_min_ = 0.1 and R_max_ = 5.1. Calcium calibration was performed using the Fura-2 calcium calibration kit (Cat. Nr. F6774, Invitrogen). Cells were identified as regions of interest (ROIs) using the free-hand selection tool of Fiji ImageJ2.0 and the mean [Ca^2+^]_i_ of each ROI for each time point was calculated and used for subsequent analysis. Data analysis was performed using custom scripts in R (see Supplementary Code). Briefly, cells with a basal mean [Ca^2+^]_i_ level higher than 200 nM before pentyl acetate application (0 ≤ Time (s) ≤ 40) or with a [Ca^2+^]_i_ standard deviation > 10 nM before pentyl acetate or VUAA1 application (0 ≤ Time (s) ≤ 40 and 300 ≤ Time (s) ≤ 340, respectively) were excluded from further analysis. Variations of the intracellular Ca^2+^ concentration (Δ[Ca^2+^]_i_) were calculated from the base level before pentyl acetate application for all analyzed cells; imaged cells coming from the same electroporation experiment (and imaged from up to 6 wells) were averaged in order to calculate the experimental time series and the responses following stimuli application. Calculated responses to stimuli were normalized to the [Ca^2+^]_i_ before each stimulus application. Statistical analysis was performed using parametric or non-parametric tests (according to the data distribution) and p-values were corrected for multiple comparisons using Holm’s correction. Software plugin and packages used are listed in the Supplementary Table 2.

Imaging experiments using an automated platform were performed on a BD Pathway 855 (BD Biosciences) controlled by the AttoVision software (version 1.6/855). Excitation at 340 (for 150 ms) and 380 nm (for 250 ms) was performed using the BD Pathway 855 settings and filters for Fura-2 with a 5 s interval between frames. Stimuli consisted of 20 µl solutions of odor at the desired concentration (see Figure 3), or 20 µl of 100 µM VUAA1 in SES. The odor solution was presented at the 10^th^ frame, while the VUAA1 solution was presented at the 30^th^ frame. After background subtraction, cells were segmented using the built-in tools of AttoVision and mean ROI intensity values were exported as text files for subsequent analysis in R (see Supplementary Code). Briefly, for experiments shown in Figure 3d-h, ROIs showing high base level fluorescence standard deviation (> 0.05 × R [340/380 nm] ratio) or a high intracellular Ca^2+^ (> 2 × R [340/380 nm] ratio) in frames 1-9 (before odor stimulus application), or showing no or very small VUAA1 responses (local max after VUAA1 application < 1.5 × baselevel R [340/380 nm] ratio) or showing an increasing monotonic VUAA1 response (i.e. no peak in the VUAA1 response before the end of the experiment) were excluded from analysis. For control experiments (Figure 3i), only ROIs with high base level fluorescence standard deviation, or with high intracellular Ca^2+^ before odor stimulus application were deleted. ROIs from wells subjected to the same treatment within the same plate were averaged (n = 1) in order to calculate the experimental time series and the responses following stimuli application. Statistical analysis was performed using parametric or non-parametric tests (according to the data distribution) and p-values were corrected for multiple comparisons using Dunnett’s correction. Software plugin and packages used are listed in the Supplementary Table 2.

### OR agonists

The following odors and agonists were used: pentyl acetate (Sigma-Aldrich, Cat. Nr. 109584, 99%), butyl acetate (Fluka, Cat. Nr. 45860, ≥99.5% (GC), ACS Reagent), methyl hexanoate (Fluka, Cat. Nr. 21599, 99.8%, analytical standard grade), propyl acetate (Sigma-Aldrich, Cat. Nr. 133108, 99%; Fluka, Cat. Nr. 40858, ≥99%, analytical standard grade), 3-methylthio-1-propanol (Sigma-Aldrich, Cat. Nr. 318396, 98%; Sigma-Aldrich, Cat. Nr. W341509, ≥98% synthetic, FG grade), hexyl acetate (Sigma-Aldrich, Cat. Nr. 108154, 99%), isobutyl acetate (Fluka, Cat. Nr. 94823, 99.8%, analytical standard grade), 2-heptanone (Sigma-Aldrich, Cat. Nr. W254401, 98% synthetic, FG grade), 3-octanol (Sigma-Aldrich, Cat. Nr. 218405, 99%), (±)-geosmin (Sigma-Adrich, Cat. Nr. UC18, ≥98% (GC)), VUAA1 (CAS Nr. 525582-84-7) was synthesized by the group “Mass Spectrometry/Proteomics” of the Max Planck Institute for Chemical Ecology (Jena, Germany). Stimuli stocks consisted of 100 mM solutions in DMSO (Sigma-Aldrich, Cat. Nr. D8418); working solutions were prepared fresh diluting 100 mM stocks in SES to the desired concentration just before the start of the experiment. Negative controls consisted of the equivalent maximum volume of DMSO used to prepare odor stimuli solutions diluted in SES.

## Results

### Targeting of insect ORs to the plasma membrane

We first tested whether the ^344^QVAPA^348^ (minimal Rho tag, abbreviated as "mRho" or "R" tag) and the ^106^VNKFSL^111^ (minimal ER export tag, abbreviated as "mER" or "E" tag) peptides from the human Rhodopsin and the HCN1 channel respectively (Figure 1a) could improve the intracellular trafficking of hOr47a to the plasma membrane in mammalian cells.

For this purpose, we performed functional imaging experiments on HEK293 cells transiently transfected with the constructs shown in Figure 1b and loaded with the calcium dye Fura-2. We stimulated the hOr47a/Orco heteromers with the Or47a agonist pentyl acetate and the Orco synthetic agonist VUAA1 (Figure 1c). We quantified the increase in intracellular free calcium with respect to the base level (∆[Ca^2+^]_i_ (nM)) (Figure 1d) and the distribution of Ca^2+^ responses (Figure 1e and Supplementary Figure 1) for each tested construct (i.e.: treatment), in order to compare the number of excited cells after presentation of the odor pentyl acetate and Orco agonist VUAA1.

HEK293 cells co-transfected with the R.E.hOr47a and Orco constructs showed significantly higher calcium responses after stimulation with 100 µM pentyl acetate and 100 µM VUAA1, compared to cells transfected with an untagged version of hOr47a (Figure 1d). Moreover, the R.E.hOr47a construct induced significantly higher calcium responses to pentyl acetate than the E.hOr47a construct bearing the mER tag alone (Figure 1d, left panel). In order to calculate and compare the number of responding cells between the tested treatments, we used the distribution of Ca^2+^ responses from HEK293 cells transfected with the empty pcDNA3.1(-) vector to calculate threshold values to classify cells as “responding” or “non responding” to an OR agonist. To do so, we calculated the mean response intensity + 2 × SD (Δ[Ca^2+^]_i_, in nM) of the top (most responsive) 0.5 percentile of the cumulative distribution of analyzed control cells (transfected with empty vector) after a pentyl acetate and VUAA1 stimulation. The resulting threshold values were 6.75 nM for pentyl acetate and 6.25 nM for VUAA1 (Supplementary Figure 1). Only 2.9 ± 1.05 % (mean ± SD) of cells transfected with hOr47a/Orco showed a ∆[Ca^2+^]_i_ ≥ 6.75 nM after a 100 µM pentyl acetate stimulation, while 21.61 ± 4.16 % of cells transfected with E.hOr47a/Orco and 30.51 ± 6.56 % of cells transfected with R.E.hOr47a/Orco reached such threshold. When comparing cell responses after a 100 µM VUAA1 stimulation, only 30.24 ± 7.21 % of cells transfected with hOr47a/Orco showed an ∆[Ca^2+^]_i_ ≥ 6.25 nM, while 61.49 ± 5.95 % and 71.76 ± 3.29 % of cells transfected with E.hOr47a/Orco and R.E.hOr47a/Orco respectively could be defined as “responding” (Figure 1e and Supplementary Figure 1). These results show that the mRho and mER tags can significantly improve the functional expression of insect ORs in mammalian cells and that the addition of mRho has a significant positive effect compared to the mER tag alone.

### Optimization of OR co-transfection in mammalian cells

To improve the chances of successful co-transfection for a tuning OR and the co-receptor Orco, we designed a new vector made of the plasmid backbone (including the *amp* resistance gene and a high-copy number *ori*) of the pCMVTNT vector and the bidirectional expression cassette of the pBI-CMV1 vector, in order to guide the expression of two genes simultaneously (see Methods). We first inserted a human codon-optimized version of *D. melanogaster* Orco (hOrco) tagged at the N-terminus with a myc tag in correspondence of the multiple cloning site 1 of the vector. As the CMV promoter can initiate expression in *E. coli* (Lewin et al., 2005) and Orco forms leaky ion channels, the unintended expression of Orco during cloning may reduce the viability of successfully transformed bacterial colonies. To avoid such inconvenience, we inserted a β-globin/IgG chimeric intron within the Orco sixth transmembrane domain. In this way, only mammalian cells can splice out the intron, leading to the production of functional Orco ion channels. The resulting plasmid, named pDmelOR, is intended to serve as a plasmid backbone to insert one of the 61 (including splice variants) tuning odorant receptors of *D. melanogaster* (Robertson et al., 2003), and the same principle can be adapted to optimize the functional expression of ORs belonging to any insect species.

In order to evaluate the performance of the bi-directional pDmelOR vector against a standard co-transfection protocol, we compared the response profile of HEK293 cells transfected with pDmelOR-R.E.hOr47a to that of cells co-transfected with an Orco and an R.E.hOr47a construct inserted in pcDNA3.1(-) each (Figure 2a-c). HEK293 cells transfected with pDmelOR-R.E.hOr47a showed a significantly higher ∆[Ca^2+^]_i_ in response to both pentyl acetate and VUAA1 stimulation, with respect to cells co-transfected with R.E.hOr47a and Orco in pcDNA3.1(-) (Figure 2d). Moreover, a significantly higher number of cells responded to pentyl acetate and after transfection with pDmelOR-R.E.hOr47a compared to cells co-transfected with R.E.hOr47a and Orco in pcDNA3.1(-) (Figure 2e). These results propose the pDmelOR vector as an efficient transfection tool for the expression of insect OR fusion constructs with higher efficacy than standard plasmid co-transfection procedures.

### Optimization of transient transfection for automated imaging platforms

Finally, we took advantage of the high level of functional expression reached combining the increased OR trafficking to the plasma membrane using the mRho and mER tags, together with the optimization of the transfection protocol due to the pDmelOR vector, to validate our method using an automated imaging platform (Figure 3).

A monoclonal stable cell line guarantees a highly homogeneous level of transgene expression within the cell population, which minimizes variability in functional assays. On the other hand, the expected variability in a population of transiently transfected cells is much higher, due to the variation in plasmid copy number between cells that results in different levels of functional expression of the gene of interest. To control for this phenomenon, we stimulated tested cells with VUAA1 100 s after the presentation of the odor stimulus, in order to use the intensity of the response to this synthetic OR agonist as a proxy of the OrX/Orco functional expression level for each cell. In this way, we could remove from downstream analyses those cells that were not expressing ORs at sufficient levels (see Methods), and we could normalize the odor response for each cell to the intensity of the VUAA1 response, thus minimizing the variability induced by the transient transfection protocol (Figure 3a-c).

Using this method we built dose-response curves for two *D. melanogaster* ORs, namely the broadly tuned receptor Or47a and the narrowly tuned Or56a, stimulated with their main agonists, pentyl acetate and geosmin respectively, and in both cases the data could be used to obtain high quality fits (Figure 3d-e). We then tested whether the tuning properties for the Or47a receptor reported in the DoOR database (Münch and Galizia, 2016) could be replicated in HEK293 cells. To do so, we selected nine odors among the known most potent agonists for Or47a (Supplementary Figure 2) and assessed their relative potency using our assay (Figure 3f). Interestingly, when ranked according to their expected potency, compared to the DMSO negative control, the odor agonists did not show a monotonic increasing pattern. Odor properties as solubility in water-based solutions may play a role in explaining such pattern. Moreover in the case of 3-octanol, the odor application significantly affected cell activity independently of Or47a expression, as cells transfected with the negative control plasmid showed a significant reduction in the cell fluorescence base level (Figure 3i). However, we further investigated whether odor purity could have influenced our results. To do so, we tested two odors (propyl acetate and 3-methylthio-1-propanol) whose relative potency differed from the expected pattern (compare Figure 3f with Supplementary Figure 2). For each odor we tested two aliquots with different chemical grade. Interestingly, while the responses to propyl acetate were not affected by the odor chemical grade (Figure 3g), the responses to 3-methylthio-1-propanol were significantly affected by this factor, indicating that 3-methylthio-1-propanol might not be an actual agonist of Or47a and the source of OR activation might originate from impurities within the extract (Figure 3h).

## Discussion

Functional expression in heterologous systems represents a key method to elucidate the function and structure of membrane protein. When confronted with the choice of which expression system to use, there are several factors to consider: from the codon usage of the gene of interest and necessary post-translational modifications (Gomes et al., 2016), to the type of downstream applications and the level of automation required. HEK293 cells represent a well-understood and very versatile choice: their transcriptome has been extensively profiled (Sultan et al., 2008; Richard et al., 2010) providing fundamental information regarding possible cross-talk between heterologous and native proteins. Furthermore, an extensive set of molecular tools has already been optimized to support protein functional expression (for an overview see Baser and van den Heuvel, 2016), and these tools are amenable to a vast array of downstream applications, from imaging to electrophysiology, that can be performed with automated and high-throughput systems (Mattiazzi Usaj et al., 2016; Obergrussberger et al., 2018).

On such a basis, we decided to implement a fast, inexpensive and versatile method for the expression of insect ORs in HEK293 cells. In order to achieve this result, we here tackled two main problems: the poor surface localization of OR proteins in mammalian cells and the limits imposed by the co-transfection of two genes (Orco and an odor-binding receptor) to obtain functional odor-gated ion channels. First, by using small peptides from the human HCN1 and Rhodopsin that are known to facilitate the release of membrane proteins from the ER and its targeting to the plasma membrane, respectively, we obtained a nearly 20-fold increase in the mean response (Figure 1d) and a 10-fold increase in the number of cells responding (Figure 1e and Supplementary Figure 1) to an odor stimulation. Then, by adopting a new high-copy number vector based on a bidirectional expression cassette (Figure 2), we significantly improved the expression efficiency of ORs, showing that three-fourths of cells were stimulated with an overall 100-fold increase in the mean ∆[Ca^2+^]_i_ in response to an odor stimulation, when compared to a standard co-transfection protocol with ORs lacking the HCN1 and Rhodopsin-derived tags (Figure 1-2 and Supplementary Figure 1). Finally, thanks to the high expression level and the possibility to use synthetic Orco agonists as internal stimulus controls to account for inter-cell variability, we showed that such system is amenable to be used with automated platforms (Figure 3) and can be consequently used for high-throughput screenings.

Although we proved the effectiveness of such system for two *D. melanogaster* ORs with different properties – a broadly tuned receptor as Or47a and the narrowly tuned Or56a – it remains to be shown how generalizable such approach is. Mammalian ORs are affected by similar problems regarding an incorrect intracellular trafficking in heterologous expression systems. Although specific classes of proteins have been shown to support the trafficking of mammalian ORs in native olfactory neurons and in heterologous systems (Saito et al., 2004; Mainland and Matsunami, 2012), conserved OR residues linked to *in silico* structural stability were shown to impact their functional expression (Ikegami et al., 2019); we cannot exclude that insect ORs are subject to similar structural constraints.

Taken together, our results show that by optimizing the intracellular trafficking and transfection conditions of insect ORs, it is possible via transient transfection to achieve expression levels in HEK293 cells that are comparable to more time and resource-demanding methods as the establishment of mammalian stable cell lines. Hence, we hope that such method can advance the study of insect ORs structure and function even in non-model organisms.

## Funding

This work was supported by the European Union’s Horizon 2020 research and innovation program under the Grant Agreement No. 662629 (F.M. and S.L.L.) and the Max Planck Society (D.W., S.S., M.K. and B.S.H.).

### Acknowledgements

The authors thank Sabine Kaltofen for help in culturing and transfecting HEK293 cells, Sascha Buchs for help with the BD Pathway 855 setup, Kerstin Weniger and Anna SpYthe with chemical profile analysis, Aleš Svatoš and Jerrit Weißflog for the synthesis of VUAA1 and Domenica Schnabelrauch for the in-house Sanger sequencing. The authors thank Antonella di Pizio (Leibniz-Institute for Food Systems Biology at the Technical University of Munich, Germany) for fruitful discussion.

## Author contributions

FM and SLL designed the study, DW and BSH contributed to the study design, SS, MK and BSH obtained funding for the project, FM and CH created the constructs, FM performed the imaging experiments, analyzed the data and wrote the first draft of the manuscript. All authors contributed to the final version of the manuscript.

## Supplementary Information

**Supplementary Table 1.**
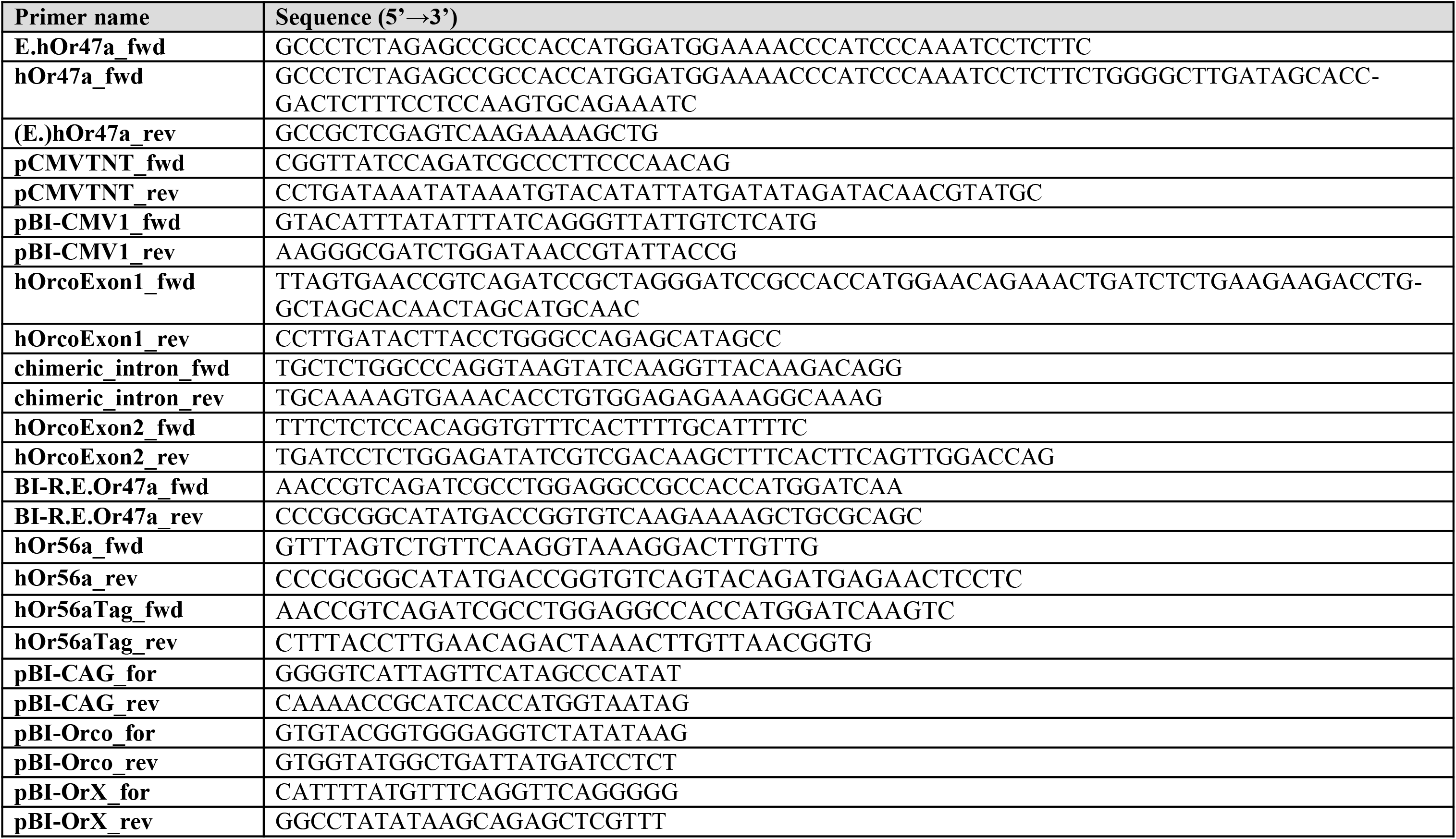
Primers used in this study.

**Supplementary Table 2.** Software plugin and packages used for data analysis.

| Platform | Plugin (Pl) or package (Pk) name | Citation |
| --- | --- | --- |
| **ImageJ** | Image Stabilizer (Pl) | K. Li, "The image stabilizer plugin for ImageJ," http://www.c-s.cmu.edu/~kangli/code/Image_Stabilizer.html, February, 2008. |
| **R** |  | R core team (2017). R: A language and environment for statistical computing. R Foundation for Statistical Computing, Vienna, Austria. URL http://www.R-project.org/. |
| **R** | ggplot2 (Pk) | H. Wickham. ggplot2: Elegant Graphics for Data Analysis (Wickham, 2016) |
| **R** | gridExtra (Pk) | (Auguie, 2017) gridExtra: Miscellaneous Functions for "Grid" Graphics. R package version 2.3.<br>https://CRAN.R-project.org/package=gridExtra |
| **R** | ggthemes (Pk) | (Arnold, 2017) ggthemes: Extra Themes, Scales and Geoms for 'ggplot2'. R package version 3.4.0.<br>https://CRAN.R-project.org/package=ggthemes |
| **R** | scales (Pk) | (Wickham, 2017) scales: Scale Functions for Visualization. R package version 0.5.0.<br>https://CRAN.R-project.org/package=scales |
| **R** | multcomp (Pk) | Torsten Hothorn, Frank Bretz and Peter Westfall (2008). Simultaneous Inference in General Parametric Models. (Hothorn et al., 2008) |
| **R** | extrafont (Pk) | (Chang, 2014) extrafont: Tools for using fonts. R package version 0.17. https://CRAN.R-project.org/package=extrafont |
| **R** | drc (Pk) | Ritz, C., Baty, F., Streibig, J. C., Gerhard, D. (2015) Dose-Response Analysis Using R (Ritz et al., 2015) |
| **R** | DescTools | Andri Signorell. 2019. DescTools: Tools for Descriptive Statistics. Version 0.99.28<br>https://cran.r-project.org/web/packages/DescTools/index.html |
| **R** | FSA (Pk) | Ogle, D.H. 2017. FSA: Fisheries Stock Analysis. R package version 0.8.17. |
| **RStudio** |  | RStudio Team (2016). RStudio: Integrated Development for R. RStudio, Inc., Boston, MA URL http://www.rstudio.com/. |

**Supplementary Figure 1.**
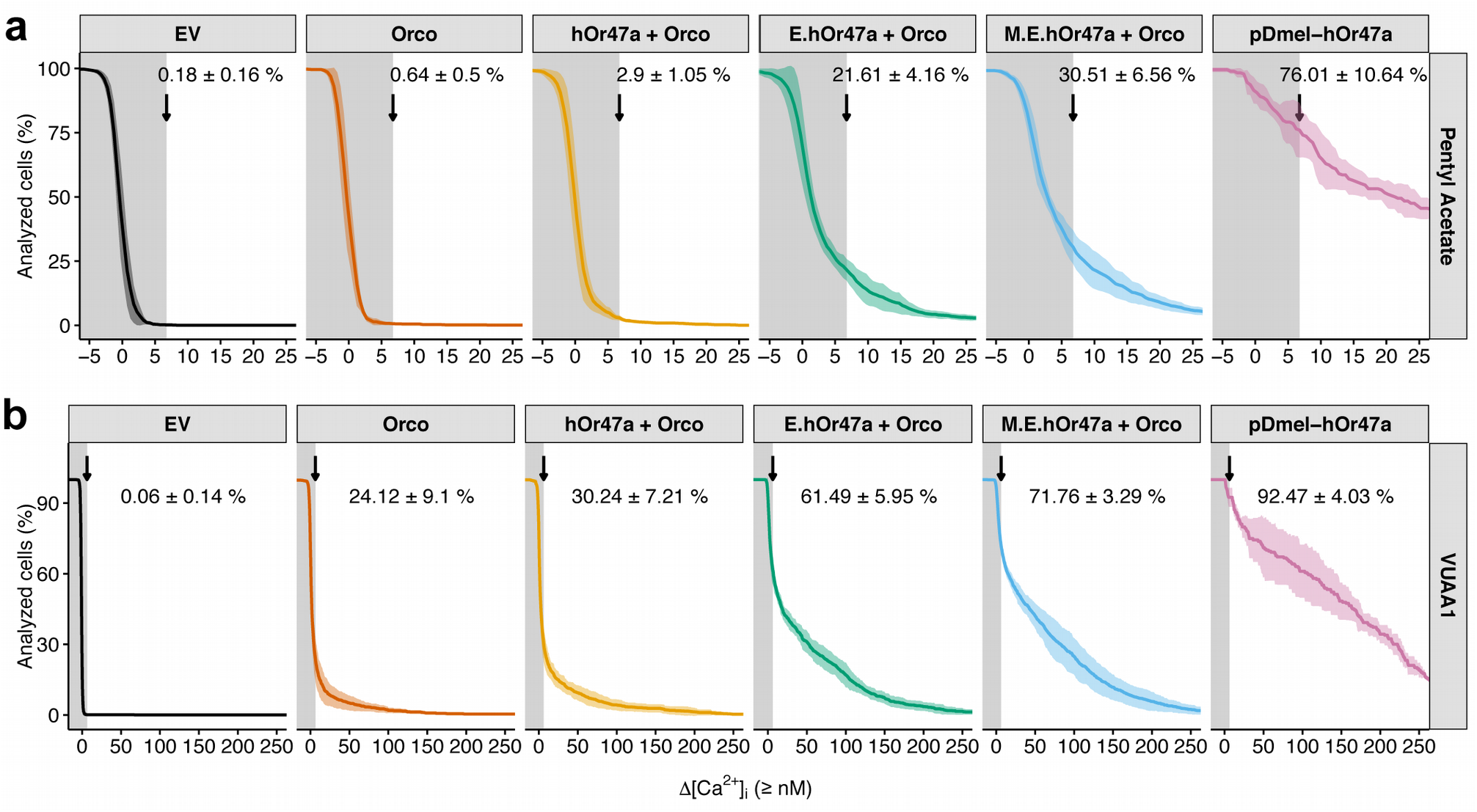
Distribution of calcium responses in transfected HEK293 cells. Distribution of the Δ[Ca^2+^]_i_ in transfected HEK293 cells following a stimulation with a 100µl of 100 µM pentyl acetate (a) or 100 µM VUAA1 (b). (a) The Δ[Ca^2+^]_i_ values were calculated for each cell 50 s after stimulation (Time = 100 s in Figure 1c and Figure 2c). Cells were identified as “responding” to the stimulus if a stimulation with pentyl acetate induced a Δ[Ca^2+^]_i_ ≥ 6.75 nM (non shaded area of the graphs). This threshold value is defined as the mean + 2 × SD response intensity (Δ[Ca^2+^]_i_, in nM) value of the top (most responsive) 0.5 percentile of the cumulative distribution of analyzed cells in the control (Empty Vector) group. (b) The Δ[Ca^2+^]_i_ values were calculated for each cell 20 s after stimulation (Time = 380 s in Figure 1c and Figure 2c). Cells were identified as “responding” to the stimulus if a stimulation with VUAA1 induced a Δ[Ca^2+^]_i_ ≥ 6.25 nM (non shaded area of the graphs). This threshold was defines in the same was as for (a). The percentage of responding cells is reported for each panel as mean ± SD. Graphs represent mean ± SD. 3 ≤ n ≤ 5 for each graph, each distribution (n = 1) is constituted by a number of cells x, with 56 ≤ x ≤ 355.

**Supplementary Figure 2.**
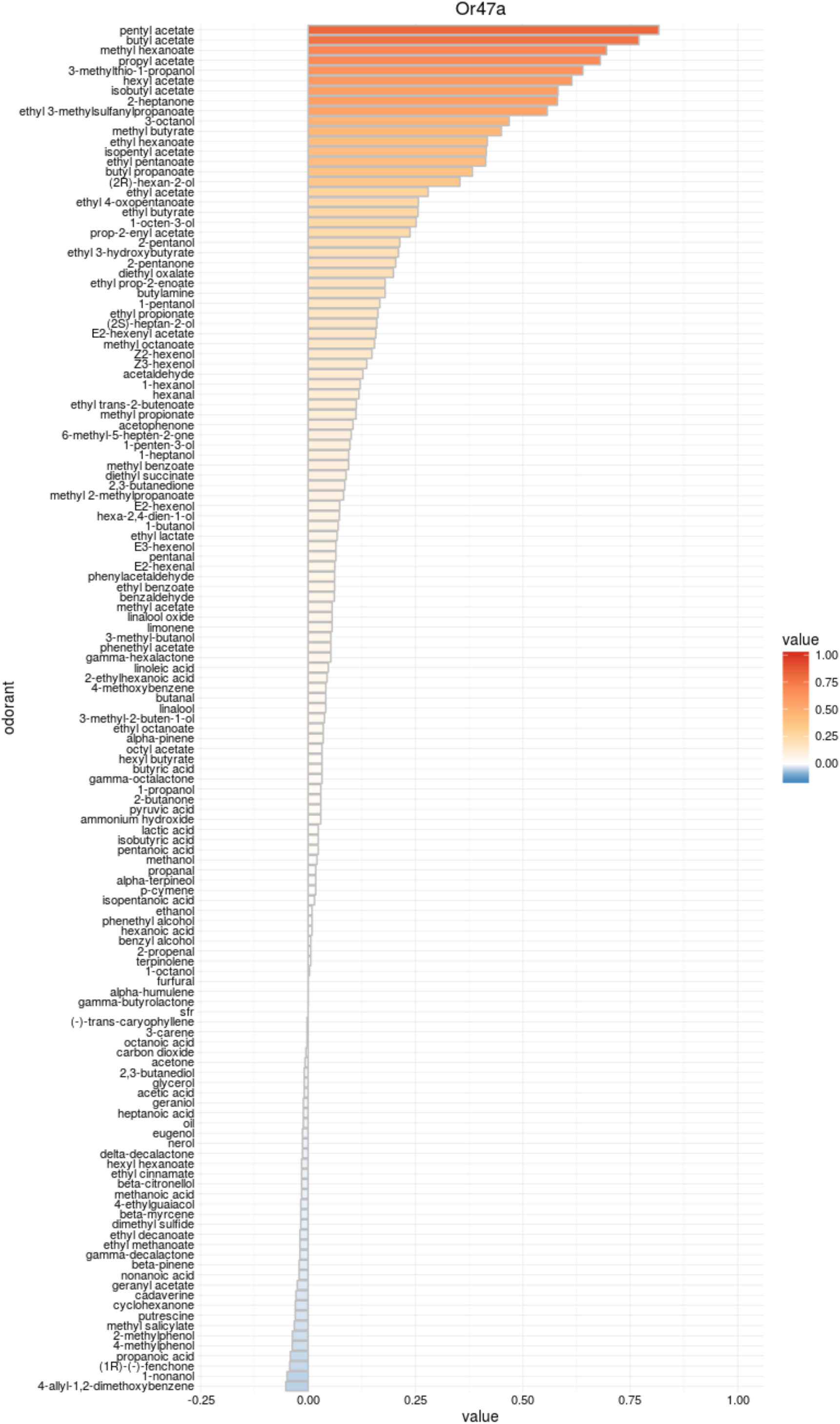
Expected odor tuning properties of *D. melanogaster* Or47a. Expected response profile for *D. melanogaster* Or47a according to the DoOR 2.0 database. Query retrieved on 24 April 2019 at http://neuro.uni-konstanz.de/DoOR/default.html.

### Supplementary Code. ImageJ and R files used for data analysis

**Fura2.ImageJ.js**: JavaScript code used to calculate [Ca^2+^]_i_ and perform cell segmentation for Figure 1-2.

**Figure_1-2.R:** R analysis for data shown in Figure 1-2 and Supplementary Figure 1.

**Figure3.R:** R analysis for data shown in Figure 3.

